# The Kinesin-3 KIF1A Functions as a Diligent Worker during Axonal Transport

**DOI:** 10.1101/2025.09.11.675527

**Authors:** Jesse Cisneros Solis, T. Lynne Blasius, Kristen J. Verhey

## Abstract

Long-distance intracellular transport is driven by motor proteins that walk along microtubule tracks. The fate of the motor protein after transport is unclear. Classically, motor proteins have been thought to function as Diligent Workers (DW) that remain attached to cargo during the entire transport event and are degraded at the end of the transport journey. In contrast, previous work suggests that kinesin-1 transport can be described by a Loose Bucket Brigade (LBB) model in which individual motor proteins participate in multiple rounds of transport. Here, we used live-cell imaging in iNeurons to test whether the kinesin-3 KIF1A functions as a DW during axonal transport. We demonstrate that the fluorescence intensity of KIF1A on particles undergoing axonal transport does not change over time, suggesting that KIF1A remains attached to its cargo for the entire transport event. We determined that KIF1A has a relatively short protein half-life, suggesting that KIF1A is degraded at the end of the journey. Moreover, protein turnover appears to be tightly controlled in iNeurons, as treating cells with proteasome inhibitors results in a cessation of KIF1A-driven transport and degradation of the KIF1A aggregates through autophagy. These results suggest that KIF1A transport fits the DW model and that KIF1A protein levels may play a role in signaling proteostatic stress in neuronal cells.

## Introduction

Intracellular transport is an important for proper cellular localization of organelles and biomolecules (i.e., cargoes) and maintenance of life. Cargo transport is especially critical in neurons where the distance between the site of synthesis (e.g., the cell body) and the protein’s final location (e.g., the presynaptic terminal) does not allow for simple diffusion to be an efficient method of transport. Active transport of cargo is achieved by molecular motor proteins that use the energy of adenosine triphosphate (ATP) hydrolysis to move along either actin filaments or microtubules of the cell’s cytoskeleton. Disruptions in axonal transport have been linked to neurodevelopmental and neurodegenerative diseases (MacGillavry and Hoogenraad 2018, Sabharwal and Koushika 2019, Sleigh et al. 2019, Aiken and Holzbaur 2021, Smith et al. 2023, Xiong and Sheng 2024).

Two types of molecular motor proteins drive long-distance transport along microtubules: cytoplasmic dynein generally transports cargo to the cell center (retrograde transport), whereas kinesin motors generally transport cargo from the cell center to the distal processes (anterograde transport). Among the kinesin superfamily, members of the kinesin-1 and kinesin-3 families play predominant roles in axonal transport. Kinesin-1 proteins (KIF5A, KIF5B, KIF5C) are involved in transport of mitochondria, lysosomes, and Piccolo-Bassoon Transport Vesicles (PTVs) to the nerve terminal, whereas the kinesin-3 KIF1A is a neuron-specific motor that primarily transports synaptic vesicle precursors (SVPs) and dense core vesicles (DCVs) (Chiba et al. 2023, Xiong and Sheng 2024).

Much work in the field has focused on uncovering the biophysical properties of kinesin-1 and kinesin-3 motor proteins and identifying adaptor proteins that specify their attachment to particular cargoes (Barlan and Gelfand 2017, Cross and Dodding 2019, Yildiz 2025). In contrast, little is known about the fate of kinesin motors upon delivery of cargo to the nerve terminal. Early studies involving the physical blockage of transport in neurons by ligation with surgical string found that kinesin-1 and kinesin-3 motor proteins accumulate on the proximal side of the blockage, with little to no motor protein observed on the distal side (Hirokawa et al. 1991, Okada et al. 1995). These studies suggested that kinesin motors function as ‘diligent workers’ which associate with cargo for the entire journey and are then degraded at the axon terminal (Figure 1A).

**Figure 1.**
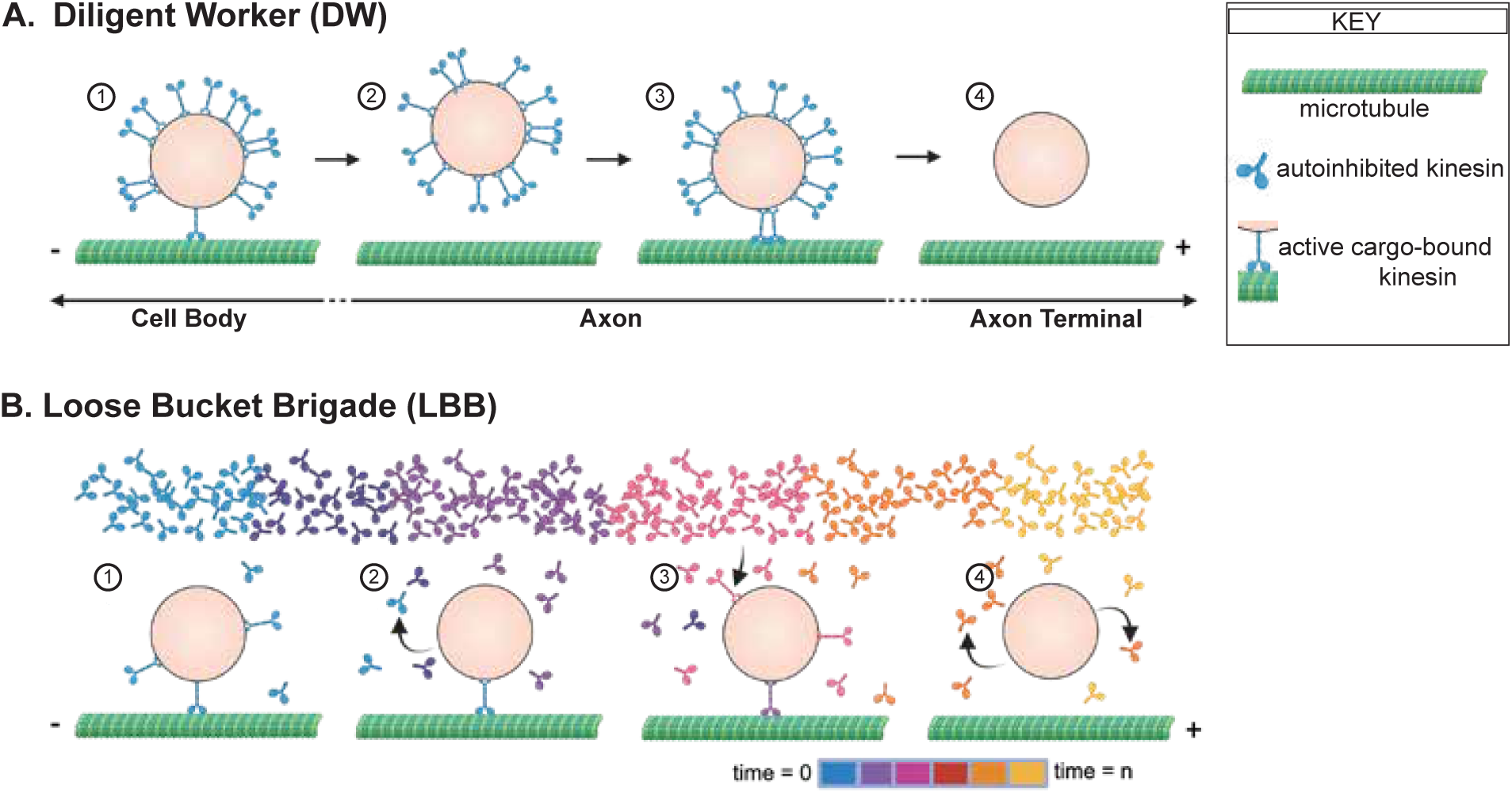
The Diligent Worker and Loose Bucket Brigade Models. Schematic of two models for kinesin-driven axonal transport. A) In the Diligent Worker (DW) model, kinesin motor proteins 1) attach to cargo in the cell body and initiate cargo transport along microtubules towards the axon terminal. Along the axon, motor-cargo complexes can 2) detach from and 3) reattach to microtubules. Upon reaching the destination, kinesin motors 4) detach from cargo and are targeted for degradation. Thus, individual motor proteins participate in a single transport event. B) In the Loose Bucket Brigade (LBB) model, there is a large pool of autoinhibited kinesin motors undergoing free diffusion throughout the cell. A few motors attach to a cargo and 1) begin microtubule-based transport towards the axon terminal. Over time, kinesin proteins 2) stochastically detach from the cargo and refold into an autoinhibited state and then different kinesin proteins 3) attach to the cargo and resume transport. Upon reaching the destination, kinesin motors 4) detach from cargo and return to autoinhibited pool. Thus, individual kinesin motor proteins are able to participate in multiple rounds of transport. Figure made with BioRender.

However, a previous study showed that kinesin-1 is not degraded upon reaching the tips of neurites in cultured neuronal cells but rather returns to the cell body via diffusion (Blasius et al. 2013). Thus, kinesin-1’s transport mode does not fit with the Diligent Worker (DW) model. Rather, through a combination of computational modeling and experimental data, kinesin-1 was proposed to function via a “Loose Bucket Brigade” (LBB) in which there is a large pool of soluble motors that transiently associate with cargoes, thereby participating in multiple rounds of transport (Figure 1B) (Blasius et al. 2013).

Whether other kinesin motor proteins function as diligent workers or in a sort of bucket brigade has not been explored. Interestingly, another study demonstrated that in *C. elegans*, UNC-104 (homolog of KIF1A) is degraded at the axon terminal via the ubiquitin-proteasome pathway (Kumar et al. 2010). This finding suggests that kinesin-3 motors may use a different transport strategy.

In this study, we used human induced pluripotent stem cells (iPSCs) differentiated into neurons and fluorescence microscopy to investigate the cargo transport strategy of KIF1A. We found that KIF1A remains attached to its cargo throughout the entire transport journey from the soma to the axon terminal, consistent with the DW model. We confirm that KIF1A has a short protein half-life, further supporting the DW model. To test whether KIF1A is degraded at the axon terminal in vertebrate neurons, we treated cells with inhibitors of the ubiquitin-proteasome pathway and found that, unexpectedly, KIF1A transport stops and KIF1A is degraded through aggrephagy in the cell body.

## Results

### KIF1A stays on its cargo during the entire journey

We set out to determine whether the kinesin-3 KIF1A behaves as a DW or whether KIF1A drives cargo transport in a LBB fashion. One difference between the two models is the stability of the motor on the cargo. In the DW model, motors remain attached to the cargo throughout the entire journey whereas in the LBB model, motors detach from cargo, undergo periods of free diffusion, and then reattach and again contribute to cargo transport (Figure 1).

To determine the stability of KIF1A on the cargo, we imaged fluorescently-labeled KIF1A motors expressed in iPSCs differentiated into neurons (iNeurons) (Gupta et al. 2018). We plated iNeurons as a low-density monolayer to minimize the overlap between neuronal processes that can make it difficult to identify and follow individual processes. We expressed full-length KIF1A tagged with mNeonGreen (KIF1A-mNG), a monomeric green fluorescent protein, and performed time-lapse imaging of neuronal processes in cells expressing low levels of KIF1A-mNG. We observed KIF1A particles traveling along the neuronal processes as well as background fluorescence from non-cargo-bound autoinhibited KIF1A motors, consistent with previous studies (Lee et al. 2003, Hung and Coleman 2016, Stucchi et al. 2018, Gan et al. 2020). To eliminate the fluorescence signal coming from the excess KIF1A motors, we implemented a bleaching protocol in which ten pre-bleach images were taken and then a region of interest (ROI) along the neuronal process was bleached at high laser power. The ROI was then imaged for an additional five minutes at one frame per second. Additional rounds of bleaching and imaging were carried out as needed, and kymographs were generated to analyze motility along the bleached axons (Figure 2A). The kymograph shows a decrease in fluorescence after each bleaching event and fluorescent KIF1A particles moving into the bleached neuronal process and traveling in the anterograde direction (Figure 2B).

**Figure 2.**
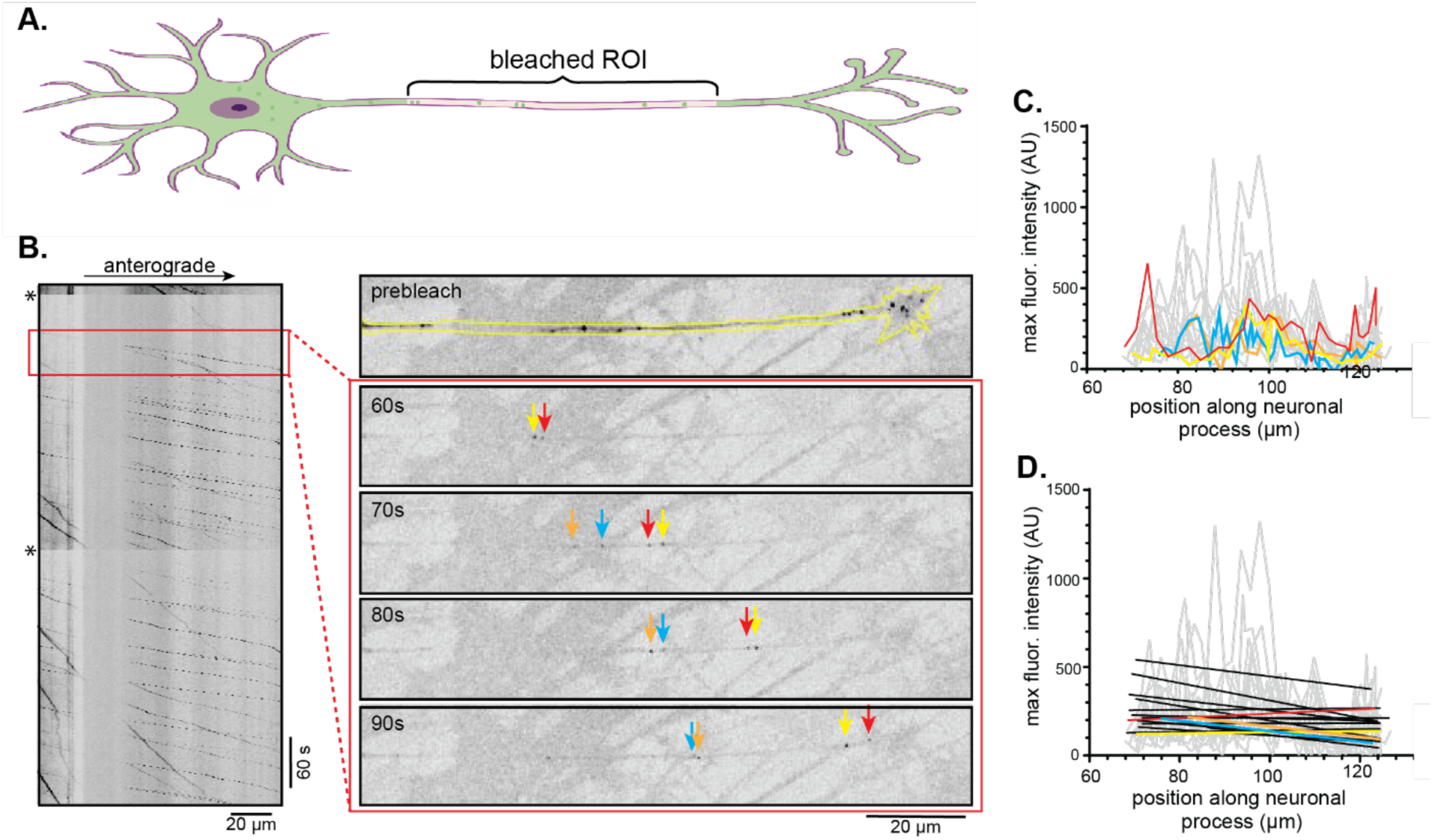
KIF1A stays attached to cargo during the transport journey. (A) Schematic of imaging assay. KIF1A-mNeGr is expressed in iNeurons, where it localizes on particles and diffusely throughout the cytoplasm. To image particles over time, a region of interest (ROI) along the neuronal process is photobleached, and then KIF1A-mNG particles that travel through this bleached region are imaged live. (B) (Left) A representative kymograph shows KIF1A particles moving anterogradely along the neuronal process after photobleaching. Asterisks indicate bleaching events. Time is on the y-axis (scale bar, 60 sec), and distance along the axon is on the x-axis (scale bar, 20 μm). (Right) Representative time frames from the red-boxed region. The yellow line in the prebleach image outlines the neuronal process, and the different colored arrows in the time frames highlight individual fluorescent KIF1A-mNG particles. (C) Graph showing the maximum fluorescence intensities of particles plotted against their position along the neuronal process (number of particles, n=16). The line colors correspond to the particles tracked in B. (D) Linear regression analysis of individual particles superimposed on the plot in (C).

If KIF1A functions as a DW, we expect the fluorescence profile of KIF1A on an individual cargo to be constant over time and distance along the axon. In contrast, if KIF1A functions in a LBB, the fluorescence profile of KIF1A on the cargo should decrease over time and distance, as there are no fluorescent freely-diffusing motors that can reattach during transit of the cargo through the bleached region. We thus measured the fluorescence intensity of individual KIF1A particles traveling anterogradely through the bleached ROI and plotted the maximum fluorescence against particle position in the neuronal process. The KIF1A fluorescence intensity appeared constant over the entire ROI, although fluctuations were observed as the neuronal process passed over and under other processes (Figure 2C, Supp Fig 1). We thus applied a linear regression analysis to the fluorescence intensities of each particle which revealed a constant fluorescence signal as the KIF1A particles traveled anterogradely through the bleached region of the axon (Figure 1D, Supp Fig 1). This result suggests that KIF1A remains bound to its cargo throughout the entire journey, supporting our hypothesis that KIF1A functions as a DW.

### Kinesin-3 KIF1A has a short protein half-life

A second difference between the DW and LBB models is the lifetime of the motor protein. In the DW model, motors are degraded at the end of a single transport event whereas in the LBB model, motors can participate in multiple rounds of transport (Figure 1). To address this, we measured the protein half-life, reasoning that if KIF1A participates in only a single round of transport, it should have a short protein half-life whereas if KIF1A participates in multiple rounds of transport in a LBB, it should have a long protein half-life similar to that reported for kinesin-1 [t_1/2_ ∼24 hr (Lee and Hollenbeck 1995, Blasius et al. 2013)].

To measure protein half-life, we treated iNeurons with inhibitors that block protein degradation by the ubiquitin-proteasome pathway. Application of the reversible proteasome inhibitor MG132 or the irreversible inhibitors lactacystin or epoxomicin resulted in an increase in KIF1A protein levels over time, consistent with degradation of KIF1A by the proteasome, whereas the kinesin-1 protein levels remained constant (Supp Figure 2A). However, we observed substantial variability between experiments in terms of how rapidly the KIF1A protein levels increased (Supp Figure 2A). We also encountered variability in the appearance of higher molecular weight bands at the 4 or 8 hr treatments (Supp Figure 2A). We note that difficulties in observing changes in soluble protein levels in neurons upon proteasome inhibition have been documented in previous studies (Ding et al. 2006, Lazarevic et al. 2011, Hakim et al. 2016, Alvarez-Castelao et al. 2020).

We thus elected to measure protein half-life by blocking new protein synthesis. In particular, the translation inhibitor cycloheximide has been widely used to measure degradation kinetics for short-lived proteins (Li et al. 2021). iNeurons were treated with cycloheximide for 0, 2, 4, 6, or 8 hours and protein levels in cell lysates were analyzed by western blot (Figure 3A). We observed a decrease in KIF1A protein levels over time and estimated the protein half-life to be approximately 4 hours (Figure 3B). In contrast, the levels of kinesin-1 protein were largely unchanged over the same time period (Figure 3B). As controls, we immunoblotted for the axonal protein superior cervical ganglia neuronal-specific 10 protein (SCG10), a neuronal-specific member of the stathmin family that is known to exhibit a short protein half-life, and GAPDH which is known to have a long protein half-life (Shin et al. 2012). Upon cycloheximide treatment, SCG10 levels rapidly decreased whereas no significant changes in the levels of GAPDH protein were observed (Figure 3). We conclude that KIF1A has a short protein half-life, which is consistent with its degradation following a single transport event in the DW model.

**Figure 3.**
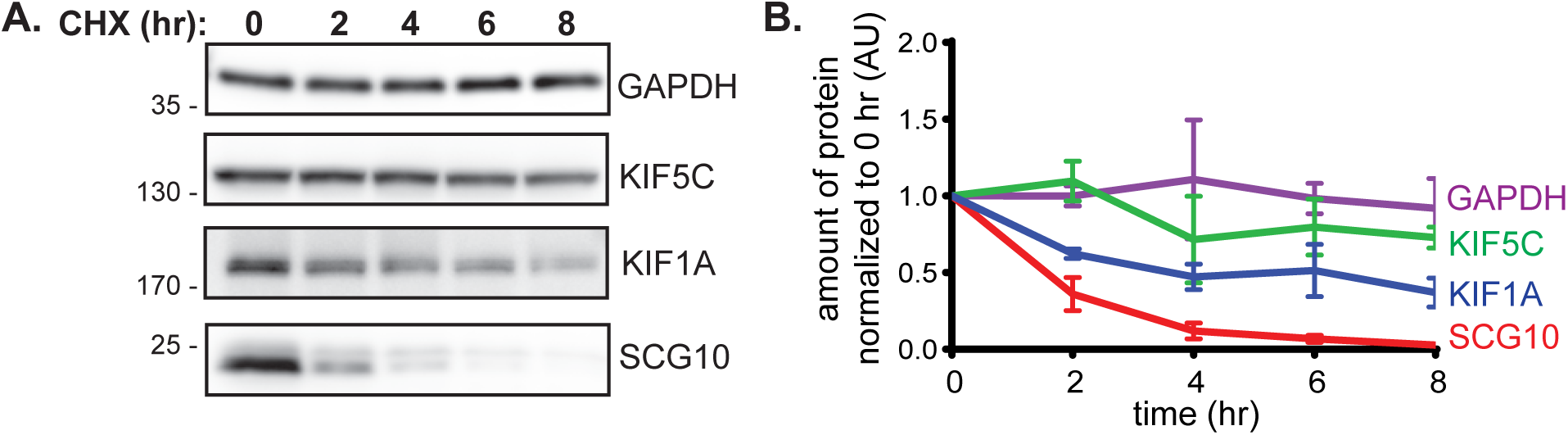
KIF1A has a short protein half-life. (A) Representative western blot of iNeurons treated with cycloheximide for various times and blotted with antibodies against the indicated proteins. (B) Quantification of the amount of protein remaining after cycloheximide treatment. The data are presented as average ± std dev from four individual experiments.

### KIF1A-driven transport stops and KIF1A accumulates in the cell body after proteasome inhibition

Another characteristic difference between the DW and LBB models is the location of motor degradation. In the DW model, kinesin motors are predicted to be degraded at the axon terminal upon completing a single transport event whereas in the LBB model, kinesin motors are proposed to participate in multiple rounds of transport and be degraded throughout the neuron (Figure 1). Therefore, we reasoned that if KIF1A functions as a DW, then it will accumulate at the growth cone when protein degradation is blocked by treating cells with a proteasome inhibitor.

To determine whether KIF1A is degraded at the axon terminal, we expressed KIF1A-mNG in iNeurons and then treated cells with the proteasome inhibitor epoxomicin for 2 hours to prevent protein degradation. The two-hour treatment was chosen to evaluate KIF1A-mNG protein levels by live imaging before the half-life of the protein is reached. We analyzed KIF1A particles localized at the growth cone in untreated and treated iNeurons by confocal microscopy (Figure 4A). In the absence of epoxomicin, we observed that the growth cone displays a spread-out morphology and contains KIF1A-mNG particles. Surprisingly, after 2 hr of epoxomicin treatment, we did not observe an accumulation of KIF1A particles or increase in KIF1A fluorescence in the growth cone (Figure 4A). In addition, the growth cone of treated iNeurons was thinner and, in some cases, retracted (Figure 4A). As an alternative approach to inhibit protein degradation, we used TAK-243 to inhibit the E1 ubiquitin-activation enzymes that carry out the first step in the ubiquitin conjugation cascade (Hyer et al. 2018). After a 2 hr treatment with TAK-243, we did not observe an accumulation of KIF1A particles or increase in KIF1A fluorescence at the growth cone (Figure 4A).

**Figure 4.**
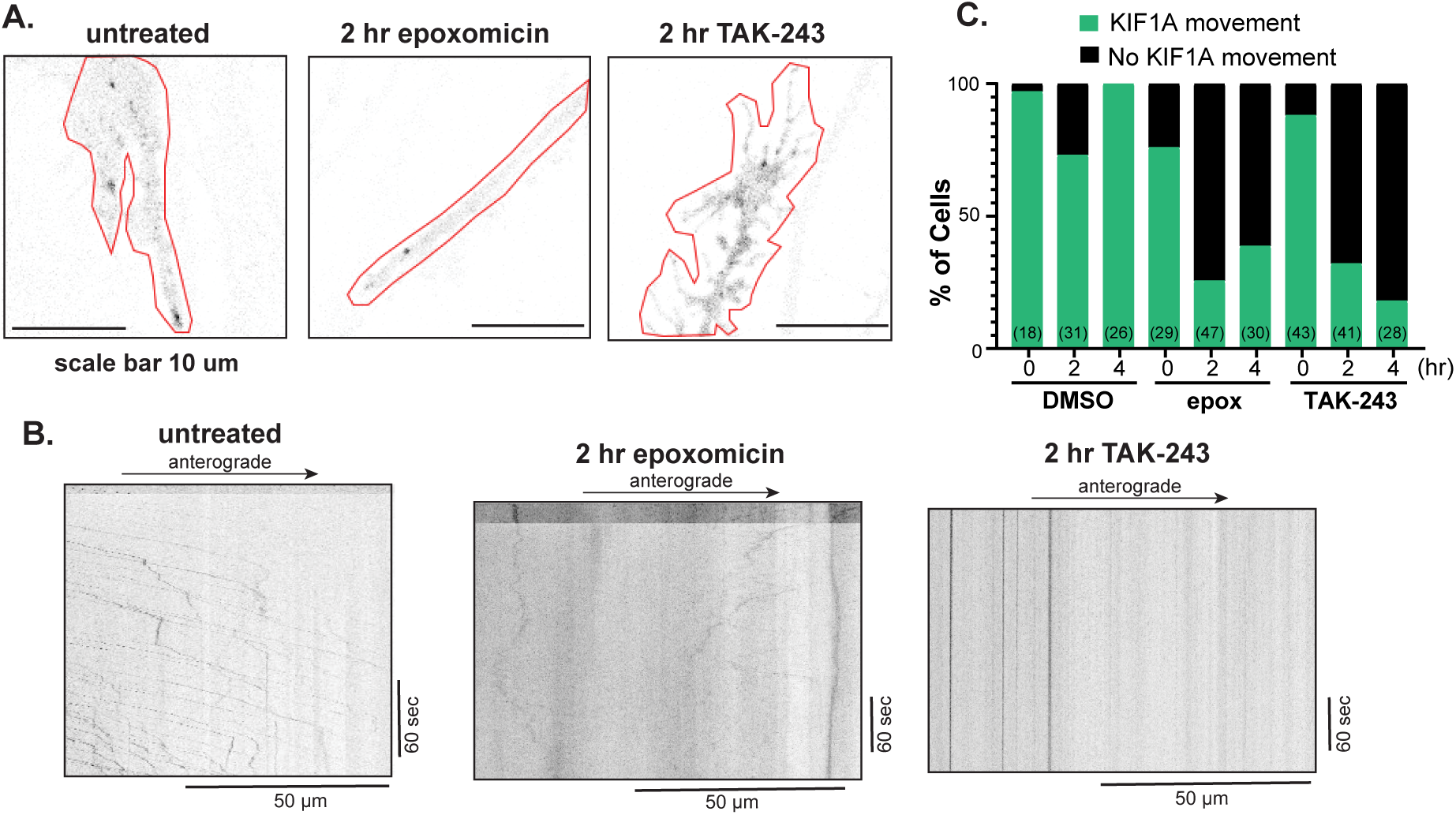
KIF1A transport decreases upon inhibition of the ubiquitin-proteasome system. (A) Representative images of KIF1A-mNG particles in the growth cones of untreated iNeurons or iNeurons treated for two hours with either 250 nM epoxomicin or 10 μM TAK-243. The red line shows the outline of the growth cone. Scale bars: 10 μm. (B) Representative kymographs of KIF1A-mNG particles traveling along the neuronal process in untreated, epoxomicin-treated, or TAK-243-treated iNeurons. Time is on the y-axis (scale bars: 60 sec), and distance is on the x-axis (scale bars: 50 μm). (C) Quantification of KIF1A-mNG particle movement along the neuronal process in untreated and treated cells. Green bars indicate cells exhibiting normal movement of KIF1A-mNG along the neuronal process. Black bars indicate cells showing no movement of KIF1A-mNG particles. The number of cells across four experiments is indicated in parentheses at the bottom of each bar.

The reduction in KIF1A particles in the growth cone after 2 hr of proteasome or E1 enzyme inhibition raises the possibility that KIF1A-driven transport is decreased after inhibition of the ubiquitin-proteasome system. To test this possibility, we performed live-cell imaging of KIF1A-mNG particles along neuronal processes of untreated or treated iNeurons. In untreated cells, we observed KIF1A-mNG traveling anterogradely along the neuronal process (Figure 4B,C). After 2 hr of epoxomicin or TAK-243 treatment, we observed little to no KIF1A-mNG particles traveling anterogradely along the neuronal process (Figure 4B,C), suggesting that inhibition of the ubiquitin-proteasome pathway results in a reduction in axonal transport.

To further investigate this, we examined the levels of KIF1A-mNG at the cell body in untreated and treated iNeurons. In untreated iNeurons, we observed numerous small KIF1A-mNG particles in the cell body (Figure 5A). In contrast, in cell treated with epoxomicin or TAK-243 for 2 hr, we observed fewer particles with higher fluorescence intensities (Figure 5A). We thus quantified the number and size of KIF1A-mNG particles following treatment with epoxomicin or TAK-243. Treatment with either inhibitor resulted in a reduction in the number of KIF1A-mNG particles in the cell body and an increase in the average size of the particles (Figure 5B,C).

**Figure 5.**
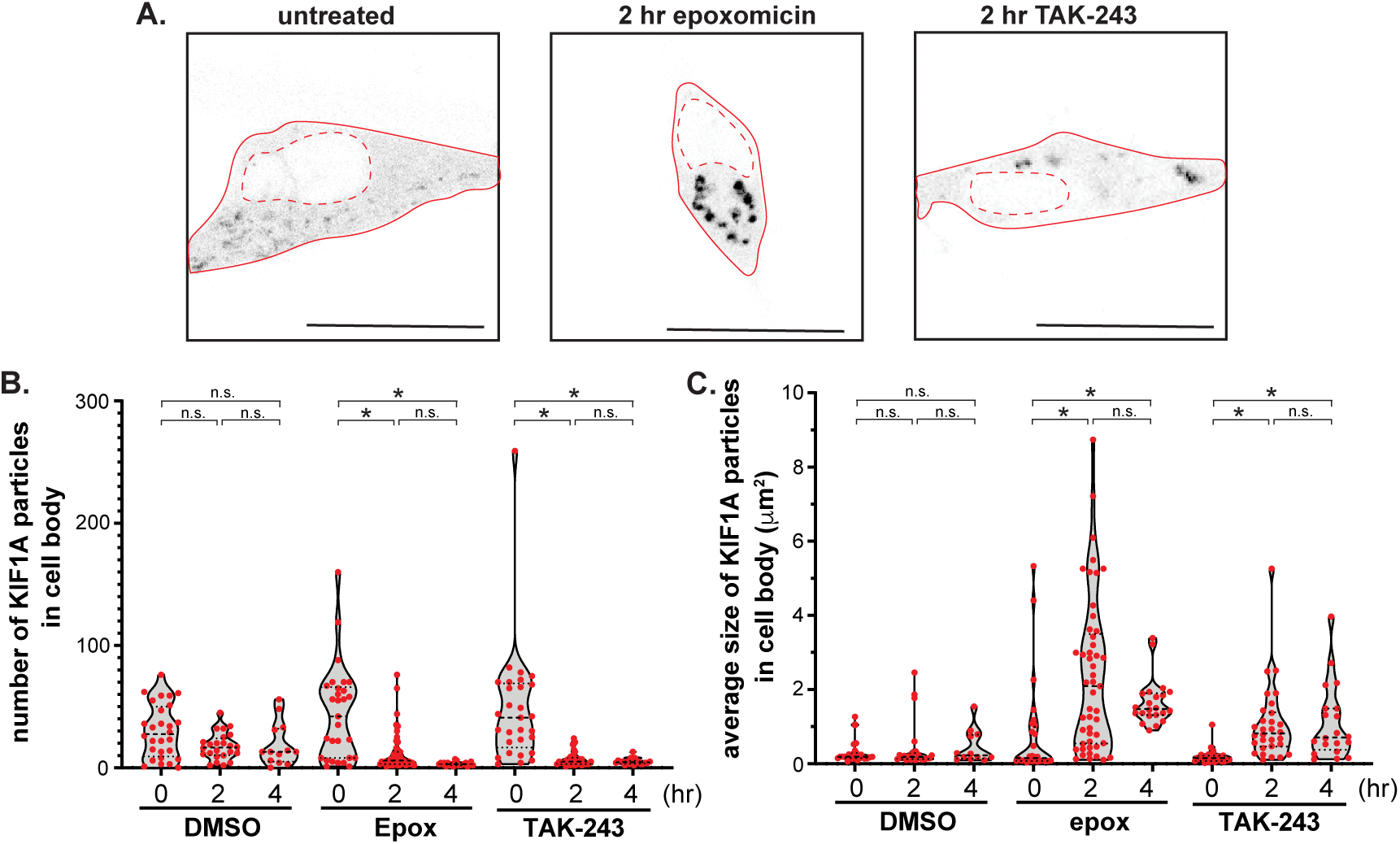
KIF1A aggregates in the cell body upon inhibition of the ubiquitin-proteasome system. (A) Representative images of KIF1A-mNG particles in the cell body of untreated iNeurons or after 2 hr of epoxomicin or TAK-243 treatment. Scale bars, 20 μm. (B) Quantification of the number of KIF1A-mNG particles in the cell body of untreated, epoxomicin, and TAK-243-treated cells. Each spot indicates the number of KIF1A particles in a single cell. n ≥ 29 cells across four or more experiments. (C) Quantification of the average size of KIF1A-mNG particles in the cell bodies of untreated, epoxomicin, and TAK-243-treated cells. Each spot indicates the average size of the KIF1A-mNG particles in a single cell. n ≥ 29 cells across more than four or more experiments. *: p<0.001, n.s.: not significant (Kruskal-Wallis test).

The increase in the size of KIF1A-mNG particles in the cell body upon inhibition of the ubiquitin-proteasome pathway could represent KIF1A-mNG molecules attached to cargo but unable to enter the neuronal process. Alternatively, the KIF1A-mNG accumulations in the cell body could represent KIF1A-mNG molecules unattached to cargo and aggregated. To distinguish between these possibilities, we examined the fate of a KIF1A cargo upon epoxomicin treatment. To do this, we expressed brain-derived neurotrophic factor (BDNF), a marker of DCVs, tagged with Halo and labeled with the dye JFX554 (BDNF-Halo^554^) in iNeurons. We analyzed the localization as well as the number and size of BDNF-Halo^554^ particles in the cell bodies of untreated and treated cells (Supp Figure 3A,B). After 2 hr of epoxomicin treatment, the size of the BDNF-Halo^554^ particles did not change, but their number in the cell body increased (Supp Figure 3C,D). The accumulated BDNF-Halo^554^ DCVs were evenly distributed through the cell body (Supp Figure 3A,B), in contrast to the KIF1A-mNG protein which accumulated near the nucleus upon epoxomicin treatment.

Taken together, these results suggest that proteasome inhibition results in a cessation of KIF1A transport and a concomitant accumulation of KIF1A protein in aggregates in the cell body, such that DCV cargoes are not transported out of the cell body.

### KIF1A accumulates in aggresomes and then in lysosomes after epoxomicin treatment

The differences in the location of KIF1A-mNG and its DCV cargo upon epoxomicin treatment suggests that KIF1A-mNG may be aggregated and targeted for degradation through aggrephagy, a selective sequestration of aggregated protein by autophagy. Autophagy was traditionally thought to be a complementary degradative pathway involving lysosome-mediated degradation but recent work suggests that impairment of the ubiquitin-proteasome pathway induces autophagy (Nam et al. 2017, Geronimo-Olvera and Massieu 2019, Finkbeiner 2020, Bauer et al. 2023, Oettinger and Yamamoto 2025).

To explore the possibility that KIF1A aggregates upon inhibition of the ubiquitin-proteasome system and that the aggregates are cleared through the autophagy pathway, we looked for colocalization of KIF1A aggregates with aggresomes and lysosomes/autolysosomes. To examine the co-localization of KIF1A with aggresomes, we expressed KIF1A-mNG in iNeurons and treated them with epoxomicin for 0, 2, or 4 hr. We then fixed and stained for aggresomes with PROTEOSTAT (Figure 6A). We observed an increase in co-localization of KIF1A-mNG with PROTEOSTAT at 2 hr of epoxomicin treatment, followed by a decrease in co-localization at 4 hrs of epoxomicin treatment (Figure 6C).

**Figure 6.**
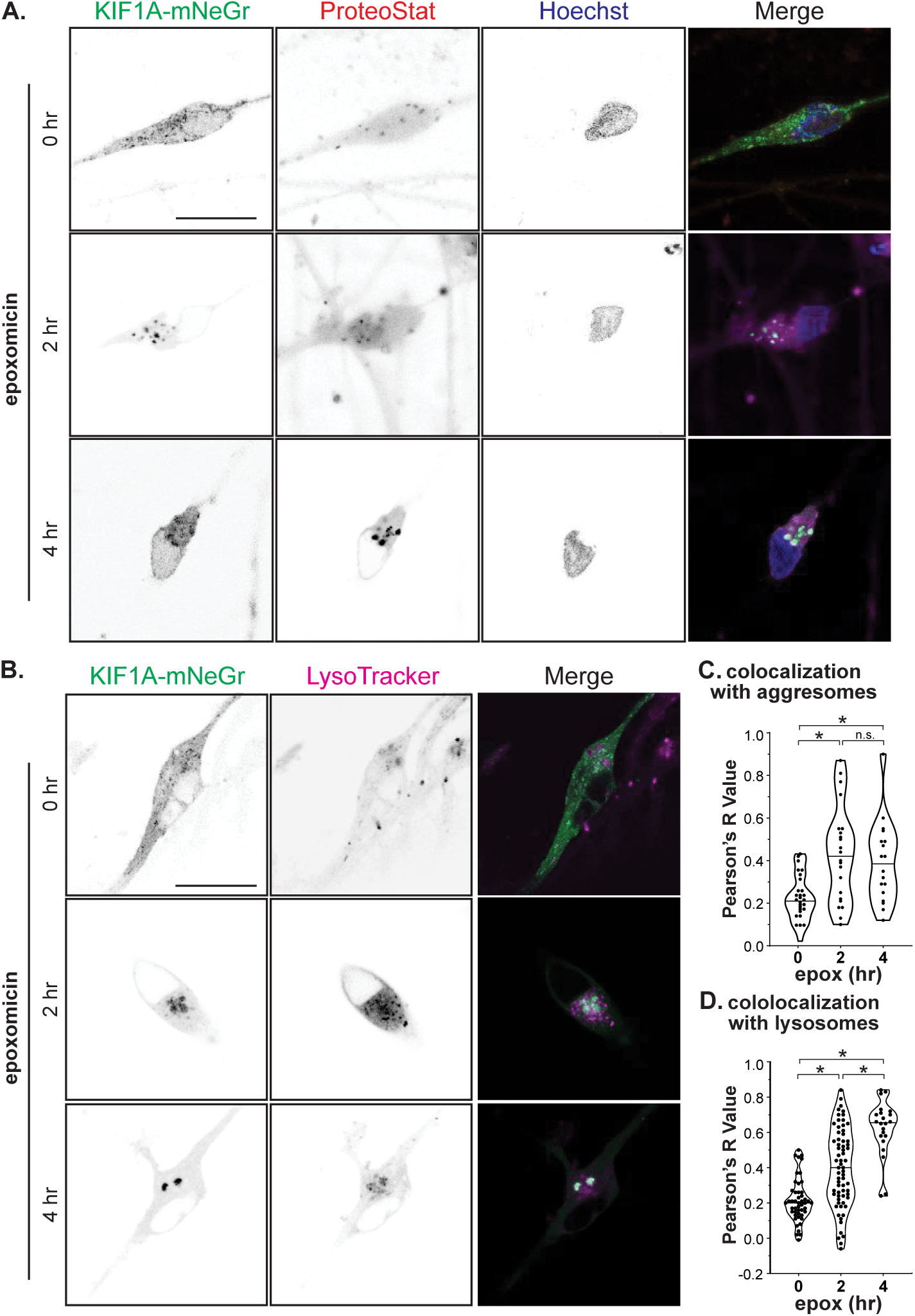
KIF1A accumulates in aggresomes and lysosomes upon proteasome inhibition. (A,B) Representative images showing the colocalization of KIF1A-mNG with A) aggresomes (ProteoStat) or B) lysosomes (Lysotracker) in the cell bodies of iNeurons treated with epoxomicin for 0, 2, or 4 hr. Scale bars: 20 μm. (C,D) Quantification of the colocalization of KIF1A-mNG with C) aggresomes or D) lysosomes over time after epoxomicin treatment. Each spot represents one cell. In C, data are from n ≥ 18 cells across one or more experiments. In D, data are from n ≥ 22 cells from one or more experiments. *: p<0.001, n.s.: not significant (Kruskal-Wallis test).

These results suggest that KIF1A-mNG is forming aggregates upon inhibition of the proteasome degradation pathway. During aggrephagy, aggregated proteins are engulfed by autophagosomes, which fuse with lysosomes to form autolysosomes, which can then degrade the aggregated protein (Lamark and Johansen 2012). We thus examined the co-localization of KIF1A with lysosomes. iNeurons expressing KIF1A-mNG were treated with epoxomicin for 0, 2, or 4 hr and stained with LysoTracker, which stains for lysosomes and autolysosomes (Figure 6B). We observed an increase in co-localization of KIF1A-mNG and LysoTracker over the epoxomicin time course (Figure 6D). These results suggest that KIF1A becomes aggregated in the cell body after epoxomicin treatment and that these aggresomes are degraded by lysosomes via the aggrephagy pathway.

## Discussion

### KIF1A transport fits the DW model

We set out to test the hypothesis that axonal transport driven by kinesin-3 KIF1A fits the DW model (Figure 1A). The DW model posits that a motor protein remains associated with its cargo throughout the entire transport journey. Indeed, using live-cell imaging and particle tracking analysis, we show that the fluorescence intensity of KIF1A-mNG on a vesicle does not change as it traverses tens to hundreds of microns along an axonal process. In addition, we observed little to no pool of freely diffusing KIF1A-mNG motors after photobleaching. These findings are consistent with previous observations of KIF1A-marked vesicles in live neurons (Lee et al. 2003, Hung and Coleman 2016, Stucchi et al. 2018, Gan et al. 2020).

A second feature of the DW model is that a motor protein is degraded upon delivery of cargo to the axon terminal. In support of this, we found that KIF1A is an intrinsically unstable protein prone to proteasomal degradation, consistent with (Huang et al. 2020). Using cycloheximide to block new protein synthesis, we estimate a ∼4 hr half-life for KIF1A in iNeurons, similar to recent work using cycloheximide to measure the half-lives of various kinesin motor proteins in differentiated PC12 cells (Huang et al. 2020). Recent work using metabolic labeling and mass spectrometry of cultured neurons have estimated the KIF1A half-life at ∼10-12 hr, but the longer estimated half-life is likely due to differences in culture conditions and/or limitations in the temporal resolution of mass spectrometry-based assays (Cohen et al. 2013, Dorrbaum et al. 2018, Ross et al. 2021). Our difficulties using proteasome inhibitors to determine protein half-life and site of degradation have been noted previously. For example, several studies have noted that treatment of cultured neurons with proteasome inhibitors did not result in the expected increase of synaptic protein levels or the accumulation of synaptic proteins at distal sites (Lazarevic et al. 2011, Hakim et al. 2016). Indeed, discerning the effects of proteasome inhibitors in neuronal cells is complicated by the finding that inhibition of the ubiquitin-proteasome system results in suppression of synaptic protein synthesis such that total protein levels can remain unaltered (Ding et al. 2006, Zhang et al. 2009, Lazarevic et al. 2011, Hakim et al. 2016, Alvarez-Castelao et al. 2020).

While these findings suggest that KIF1A motors are degraded after a single round of transport, we were unable to determine whether KIF1A is degraded at the axon terminal as treatment of iNeurons with inhibitors of the ubiquitin-proteasome pathway resulted in a cessation of KIF1A-mediated transport. However, several lines of evidence support the hypothesis that KIF1A motors are degraded upon the completion of their transport journey rather than being recycled for further use. First, in *C. elegans* mechanosensory neurons, genetic inactivation of the ubiquitin-proteasome pathway results in the accumulation of UNC-104 protein at the axon terminal (Kumar et al. 2010). Second, using compartmentalized neuronal cultures to separate axons from cell bodies, Huang and colleagues found that the proteasomal degradation of KIF1A protein occurs in axons (Huang et al. 2020). Taken together, these results suggest that KIF1A transport fits with the DW model.

### Cargo transport via the DW and LBB models

We consider the DW and LBB models to be the extreme models for how kinesins associate with their cargoes to drive axonal transport (Figure 1). The DW model provides a clear advantage for transporting cargo over extended distances, such as the axon of a neuronal cell, as multiple motors remain bound to the cargo. In the LBB model, the stochastic attachment of kinesin-1 motors improves the efficiency of cargo transport (Conway et al. 2012), yet the LBB model is limited over extended distances since the efficiency of motor diffusion back to the cell body decreases with increasing axonal length (Blasius et al. 2013).

Autoinhibition of non-cargo-bound motors is a critical component of the LBB model, yet it seems likely that autoinhibition is also important for regulation of kinesins in the DW model. We hypothesize that KIF1A autoinhibition is critical in the cell body before binding cargo and at the axon terminal after delivery of cargo. In support of this, mutations that impair the ability of UNC-104 to bind to synaptic vesicle cargoes result in a loss of UNC-104 protein, suggesting that non-cargo bound motors are degraded (Kumar et al. 2010). In addition, mutations in UNC-104 and KIF1A that abolish autoinhibition (i.e. constitutively active motors) result in higher levels of KIF1A protein at the axon terminal (Niwa et al. 2016, Chiba et al. 2019, Cong et al. 2021), suggesting that constitutively active motors are not degraded.

Intraflagellar transport (IFT) in eukaryotic cilia appears to utilize an intermediate strategy with features of both models. In IFT, kinesin-2 motors stay associated with their cargo for the entire journey and then diffuse back to the base of the cilium. Diffusion is not limiting in this system due to the shorter length of the cilium (∼10μm in *Chlamydomonas*) and the return of kinesin-2 motors via diffusion has been suggested to underlie the length-control mechanism of this organelle (Engel et al. 2009, Ludington et al. 2013, Chien et al. 2017, Hendel et al. 2018, Li et al. 2020, Ma et al. 2020).

### Inhibitors of the ubiquitin-proteasome system result in stoppage of axonal transport and aggregation of KIF1A in the cell body

We demonstrate that treatment of cultured iNeurons with inhibitors of the ubiquitin-proteasome system results in cessation of KIF1A-mediated axonal transport within 2 hr. We also find that inhibition of the ubiquitin-proteasome system results in the localization of KIF1A aggregates in the cell body. These findings extend recent work demonstrating that proteasome inhibitors result in the accumulation of proteasomes, synaptic proteins, and reporter proteins in aggresomes and autophagosomes in the cell body (Laser et al. 2003, Shen et al. 2013, Staff et al. 2013, Ale et al. 2015, Choi et al. 2020, Blumenstock et al. 2021). These results can also explain why we encountered difficulties in measuring changes in soluble KIF1A protein by western blotting.

Further work is needed to understand how KIF1A aggregates localize in the cell body. One possibility is that upon inhibition of proteasome activity, KIF1A is ubiquitinated at the axon terminal and returned to the cell body by retrograde transport. While it is known that KIF1A moves in both anterograde and retrograde directions in axons, the ubiquitinated or other state of KIF1A undergoing retrograde transport is not known. Although we did not observe an increase in retrograde transport of KIF1A-mNG at 2 hr of proteasome inhibition, imaging at earlier time points after proteasome inhibition may reveal changes in retrograde transport rates.

A second possibility is that upon inhibition of proteasome activity, KIF1A is ubiquitinated at the axon terminal where it aggregates and is engulfed by autophagosomes that undergo retrograde transport back to the cell body. This possibility is supported by studies showing that under normal cellular conditions, autophagosomes form at axon terminals and undergo dynein-dependent transport to the cell body for lysosomal clearance (Kargbo-Hill and Colon-Ramos 2020, Cai and Ganesan 2022, Sidibe et al. 2022, Lienard et al. 2024).

A third possibility is that upon inhibition of proteasome activity, there is a proteostatic stress response that stops anterograde transport and targets KIF1A for aggrephagy. This possibility is supported by recent work showing that when the ubiquitin-proteasome system is blocked, misfolded proteins and inhibited proteasomes accumulate in aggresomes in the cell body and are cleared by aggrephagy (Shen et al. 2013, Kageyama et al. 2014, Choi et al. 2020, Mukherjee et al. 2021). It may be that the relative levels of ubiquitinated and unubiquitinated proteins in the axon terminal are sensed in some way to initiate retrograde signaling pathway. It is also possible that the levels of KIF1A itself or one of its synaptic protein cargoes are sensed by the cell. In either case, it is tempting to speculate that a retrograde signaling pathway is initiated, perhaps similar to the axonal injury response pathway involving the dual leucine zipper kinase [DLK, Wallenda (Wnd) in flies]. Wnd/DLK activation is essential for injury-induced retrograde signaling that results in differential gene expression that reduces the synthesis of synaptic proteins, thereby lowering the axonal transport load (Asghari Adib et al. 2018, Jin and Zheng 2019, Smith et al. 2020). Interestingly, recent work showed that the absence of UNC-104 protein activates the Wnd/DLK signaling pathway whereas mutations that impair kinesin-1 or dynein transport do not (Xiong et al. 2010, Li et al. 2017). Our work suggests that an excess of KIF1A protein at the axon terminal may also serve as a signal of protestatic stress under conditions when the degradation pathways are overwhelmed. The sensing of efficient proteostasis is essential to neuronal health and dysregulation of the proteostasis network can lead to neurodegenerative disease (Di Domenico and Lanzillotta 2022, Kinger et al. 2024, Selvaraj et al. 2025).

## Materials and Methods

### Cell Culture

An induced pluripotent stem cell (iPSC) line, HITNC, was derived from commercially available human foreskin fibroblasts (MTI Global Systems GSC-3002) and reprogrammed using episomal plasmids as described (Okita et al. 2011). iNeurons were derived from HITNC by stable integration of an expression vector (pMT-TRE-hNGN2) for tetracycline regulated expression (TRE) of human neurogenin-2 (hNEUROG2) using Tol2 recombinase as described (Gupta et al. 2018).

iPSCs were maintained on 6-well plates coated with Geltrex LDEV-free reduced growth factor basement membrane (Matrigel, ThermoFisher A1413202) in Gibco Essential 8 (E8) Flex Medium (ThermoFisher A2858501) and passaged every 4– 5 days with EDTA (Lonza BMA51201). Coated plates were prepared by incubation in 166μg/mL Matrigel in DMEM/F12 (Gibco 14175095) for 1 h at room temperature on a rocker. An equal volume of DPBS (Invitrogen 14190144) was added and the plates were stored at 37°C.

iNeurons were induced from iPSCs by plating in E8 media with 10 mM Rock inhibitor Y-27632 (Tocris 1254). The following day, the cells were treated with 0.5 mg/mL doxycycline (Sigma D9891) to induce NEUROG2 gene expression. On day 3 when the majority of cells ceased proliferation, the cells were passaged onto laminin-coated dishes in 3N differentiation media [1:1 DMEM/F-12:Neurobasal (Gibco 21103049), 1 mM Glutamax (Gibco 35050061), 5 mg/mL insulin (Sigma I9278), N2 supplement (Gibco 17502048), B27 supplement (Gibco 17504044), 1 mM Non Essential Amino Acids (Gibco 11140050), Penn/Strep (Gibco 15140122), 2 µl β-mercaptoethanol (Sigma M3148), 0.5 mg/mL doxycycline (Sigma D9891)]. Half of the media was replaced daily with fresh 3N media until analysis. Alternatively, on day 3, the cells were resuspended in freezing medium (60% DMEM/F12, 30% KOSR (Invitrogen 10828-010), and 10% DMSO (Sigma D2650) and stored in liquid nitrogen. When needed, the cells were thawed and plated directly on laminin-coated glass-bottom dishes.

To prepare laminin-coated dishes, 35 mm glass-bottom dishes (MatTek P35G-1.5-14-C) were acid-washed by incubation in HCl for 15 min at room temperature. The dishes were washed 2x with 3 mL DPBS and then 3x with 4 mL Corning cell culture water (25-055-CM) and then incubated overnight at room temperature in 2 mL PEI/borate [110 mg of PEI (Sigma 408727)/50 mL borate buffer (ThermoScientific 28384)]. The following day, the dishes were washed with 3x with 4 mL Corning cell culture water and allowed to air dry for ∼40 min with the lids off before addition of 3.3 μg of mouse laminin per 35 mm dish [Sigma L2020, diluted in Hank’s Balance Salt Solution (Gibco 14175095)] and incubation for 2-4 hr at 37°C.

All human iPSC cell culture experiments were approved by the University of Michigan Human Pluripotent Stem Cell Research Oversight Committee (HPRSCRO record #1118). All cell lines are checked annually for mycoplasma contamination.

### Plasmids and Transfection

To construct pBa-KIF1A-mNG for expression of full-length KIF1A driven by the beta actin promoter, the rat KIF1A coding sequence (Uniprot F1M4A4) from pKIF1A-mCit-N1 (Hammond et al. 2009) and the mNG fluorescent protein from pmNeonGreen-N1 were inserted into pBa-KIF1A(1-396)-GFP vector [gift of Gary Banker & Marvin Bentley, Addgene plasmid #45058 (Jacobson et al. 2006)] using convenient restriction sites. pBa-BDNF-Halo was a gift from Marvin Bentley [Addgene plasmid # 162718 (Frank et al. 2020)]. iNeurons were transfected on differentiation day 7 with 500 ng plasmid DNA and Lipofectamine LTX with PLUS reagent (ThermoFisher 15338100) per manufacturer’s instructions and the cells were imaged 24 hours after transfection.

### Live-cell Imaging

For live-cell imaging in Figure 2, images were acquired using a Nikon N-SIM Superresolution and A1R laser scanning confocal microscope equipped with a plan apo lambda 60x/NA 1.4 oil-immersion objective. A stage-top live-cell chamber was used to maintain a temperature of 37°C and an atmosphere of 5% CO_2_. The media was replaced with prewarmed L15 media (Gibco 21083027). KIF1A-mNG images were captured at 5% laser power, with 488 nm excitation, using a filter cube of 525/50 at a rate of 1 frame per second (fps) for 10 seconds. Photobleaching of a Region of Interest (ROI) along the axon of a KIF1A-mNG-expressing cell was conducted using a 488 nm laser at 100% intensity for 10 seconds. Following the bleaching, imaging resumed with 5% laser power and 488 nm excitation at 1 fps for an additional 5 minutes. Only particles moving in the anterograde direction within the neuronal process were tracked using the Fiji plug-in “Manual Tracking,” with the centering option set to local maximum and a searching square size of 5 pixels. Kymograph and tracking were generated using ImageJ 2.14 and fluorescence intensity over time plot and linear regression analysis were carried out using GraphPad Prism 10.2.

For live-cell imaging in the absence and presence of proteasome inhibitors (Figures 4-6), iNeurons were treated with 250 nM Epoxomicin (UBPBio F1400) or 1 uM TAK-243 (MedChemExpress HY-100487) for 0, 2 or 4 hr. For iNeurons transfected with BDNF-Halo, the JFX554 ligand (Promega HT1030) bwas added 30 min before imaging. The media was replaced with prewarmed L15 media and the cells were imaged using a Zeiss Airyscan2 laser scanning confocal microscope with a plan apo 63x/NA 1.4 oil immersion objective. The microscope stage was equipped with a live-cell chamber to maintain a temperature of 37°C and 5% CO_2_. KIF1A-mNG images were captured at a laser power of 2.5-5% with 488 nm excitation. BDNF-Halo images were obtained at a laser power of 5% with 561 nm excitation. The size and number of particles in the cell bodies were analyzed using the “Analyze Particle” plug-in in Fiji using parameters Gaussian blur (sigma: 0.50), convolution, and thresholding. The resulting plots were generated using GraphPad Prism 10.2.

For colocalization of KIF1A-mNG with lysosomes, iNeurons were treated with 100 nM LysoTracker Deep Red (Invitrogen L12492) for 2 hr before imaging. The media was replaced with prewarmed L15 media and the iNeurons were imaged live on a Zeiss LSM 980 Airyscan 2 laser scanning confocal microscope with a plan apo 63x/NA 1.4 oil immersion objective. The stage was equipped with a live-cell chamber to maintain 37 °C and 5% CO_2_. Snapshots of KIF1A-mNeonGreen were taken at 2.5-5% laser power of 488 nm excitation and of LysoTracker at 0.5% laser power of 639 nm excitation. The size and number of particles in the cell bodies were analyzed using the “Analyze Particle” plug-in in Fiji (ImageJ 2.14) using parameters Gaussian blur (sigma: 0.50), convolution, and thresholding. Colocalization was calculated using using Pearson’s correlation coefficient (R) with the “CoLoc2” plug-in and the plots were generated using GraphPad Prism 10.2.

### Fixed Cell Imaging

For colocalization of KIF1A-mNG with the aggresome, iNeurons were fixed in 3.7% paraformaldehyde and 4% sucrose in PBS at room temperature for 30 min and then permeabilized using 0.05% Triton X-100 and 4% sucrose in PBS on ice for 30 min. Then the cells were stained with PROTEOSTAT Aggresome Detection Reagent (Enzo Life Sciences ENZ-51035 (Shen et al. 2011)) prepared according to the manufacturer’s protocol. Snapshots of KIF1A-mNeonGreen were taken at 2.5-5% laser power at 488 nm excitation and of Proteostat at 5% laser power at 561 nm excitation. The size and number of particles in the cell bodies were analyzed using the “Analyze Particle” plug-in in Fiji using parameters Gaussian blur (sigma: 0.50), convolution, and thresholding. Colocalization was quantified using Pearson’s correlation coefficient (R) with the “CoLoc2” plug-in in Fiji. Graphs were generated and statistical analyses were applied using GraphPad Prism 10.2.

### Western Blot

iNeurons at differentiation day 8 on a Matrigel-coated 6-well plate were treated for 0, 2, 4, or 8 hours with 100 μg/mL cycloheximide (VWR 97064-724), 250 nM epoxomicin (UBPBio F1400), lactacystin (Cayman Chemical 70980), or MG-132 (Sigma M7449). Cell lysates were prepared by trypsinizing with 0.05% Trypsin–EDTA (Gibco 05300054), pelleting the cells at 1500 × g for 5 min at 4°C, and aspirating the supernatant. The cell pellet was resuspended in growth medium and then pelleted again. The cell pellets were resuspended in lysis buffer (25 mM HEPES, 115 mM KOAc, 5 mM NaOAc, 5 mM MgCl_2_, 0.5 mM EGTA, 1% Triton X-100 v/v)

and spun at 18,213 × g for 10 min at 4°C. Supernatants were collected and stored at -20°C. 1/20th of a 35 mm well of iPSCswere loaded per well and the proteins were separated by SDS-PAGE and transferred to a nitrocellulose membrane. Western blot was carried out with antibodies against KIF1A (BD Transduction Laboratories 612094), kinesin-1 (clone H2, Millipore Sigma MAB1614), GAPDH (clone EPR16891 AbCam Ab181602), and SCG10 (Novus Biologicals NBP1-49461). Protein levels were quantified using ImageJ 2.14 and plotted using GraphPad Prism 10.2.

---

**Supplemental Figure 1.**
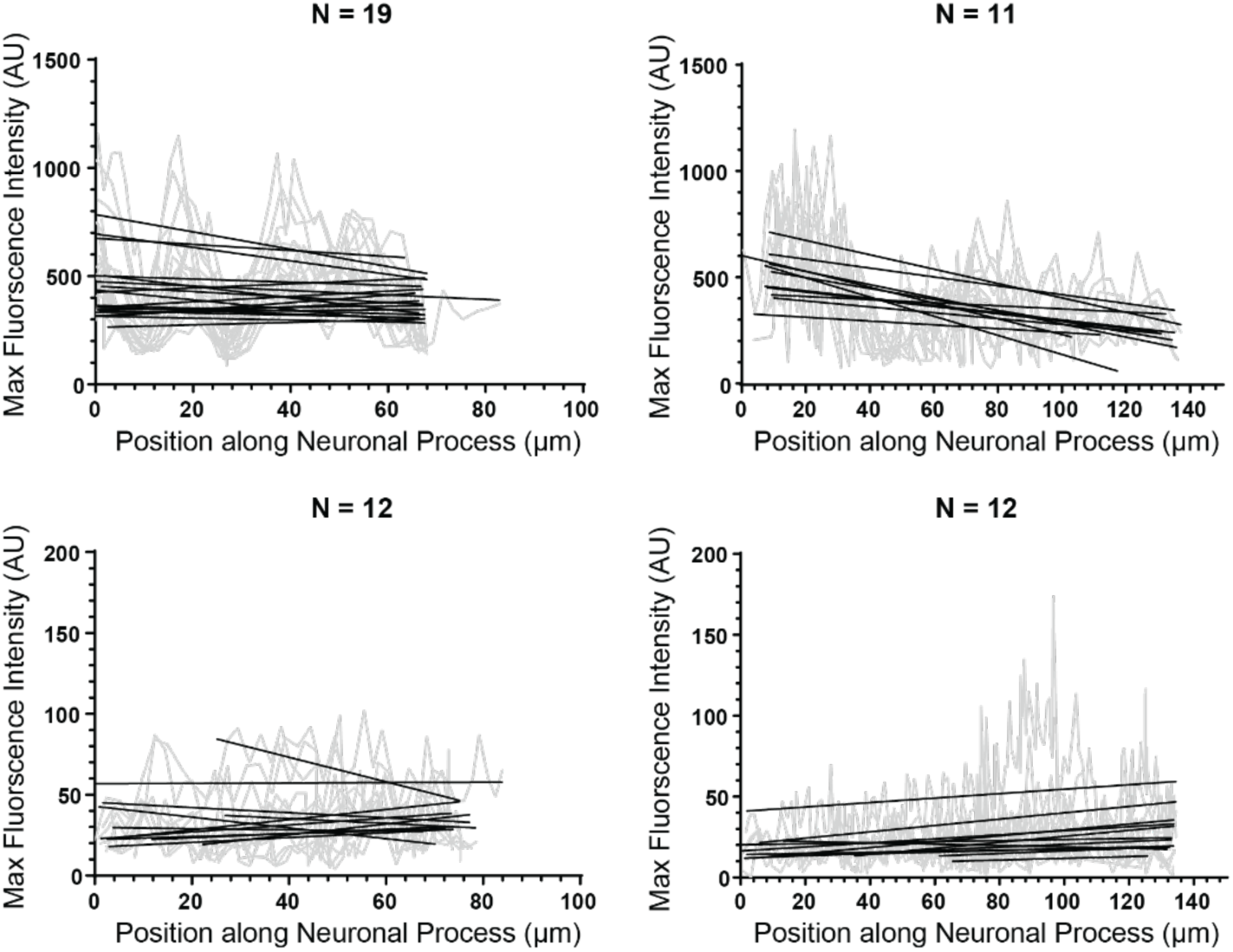
KIF1A stays attached to cargo during transport. Additional examples of the maximum fluorescence intensities of KIF1A-mNG particles plotted against their position along the neuronal process, with linear regression analysis of individual particles superimposed. Each graph represents a single cell. The N value above each graph indicates the number of particles.

**Supplemental Figure 2.**
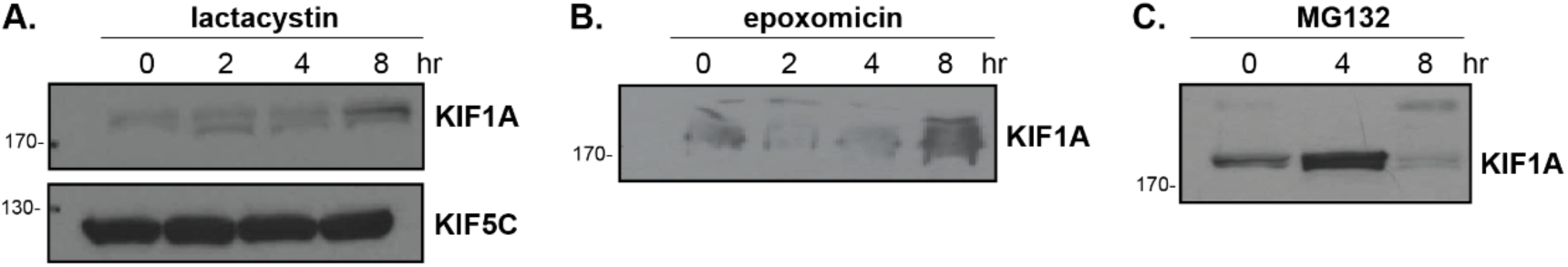
KIF1A has a short half-life. Representative western blots of soluble proteins in cell lysates of iNeurons treated with the proteasome inhibitors A) lactacystin, B) epoxomicin or C) MG132 for the indicated times. Nitrocellulaose membranes were blotted with antibodies against the kinesin-3 KIF1A or the kinesin-1 KIF5C.

**Supplemental Figure 3.**
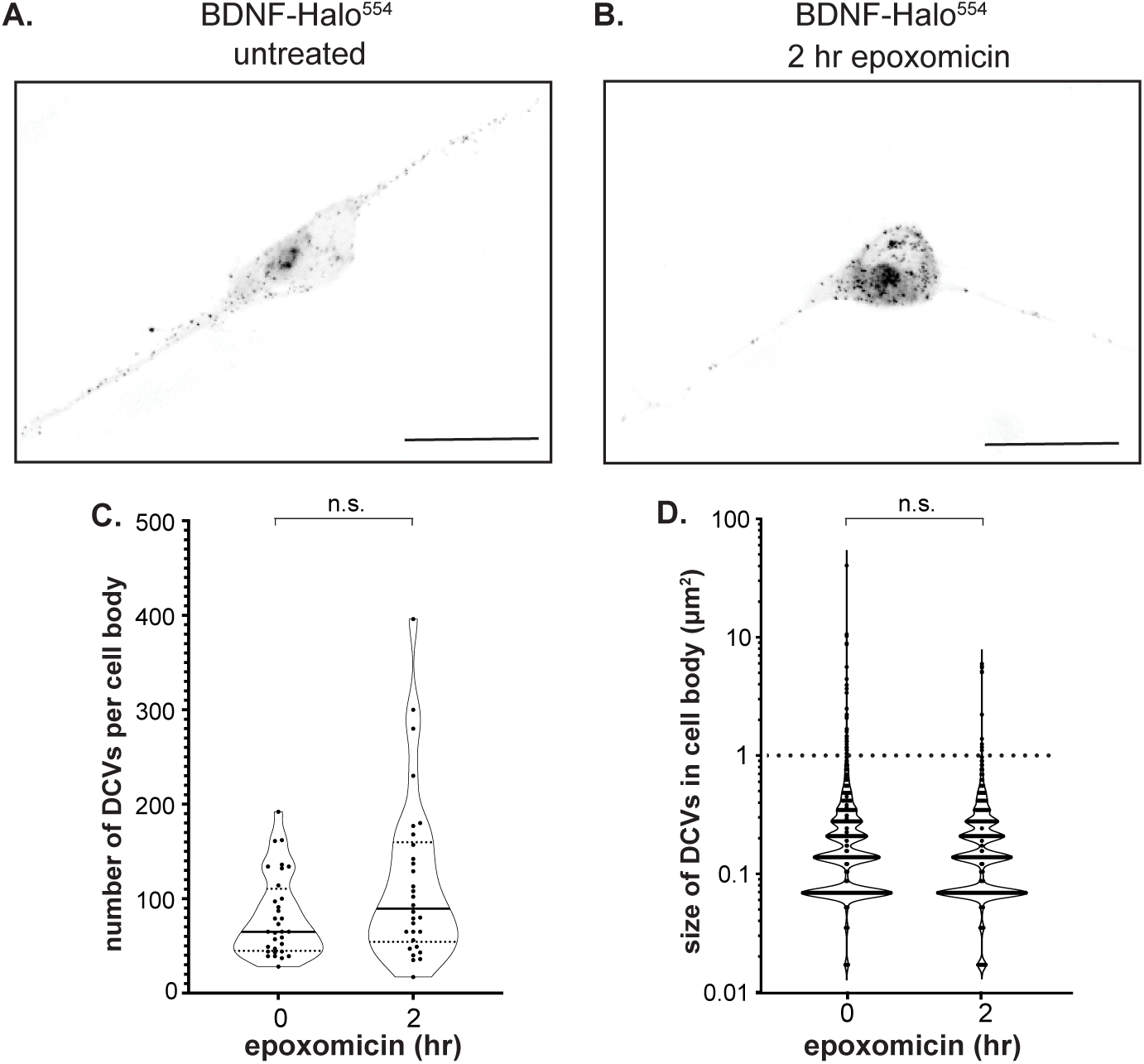
DCV transport stops after proteasome inhibition. (A, B) Representative images of BDNF-Halo^554^ particles in the cell bodies of (A) untreated cells or (B) cells treated with epoxomicin for 2 hr. Scale bars: 20 μm. (C,D) Quantification of the C) number and D) size of BDNF-Halo^554^ particles in the cell bodies of untreated versus epoxomicin-treated cells. Each spot represents a single cell. n ≥ 30 cells across 4 experiments, n.s.: not significant (Kruskal-Wallis test).

## References

Aiken, J. and Holzbaur, E. L. F. (2021). Cytoskeletal regulation guides neuronal trafficking to effectively supply the synapse. Curr Biol, 31, R633–R650.

Ale, A., Bruna, J., Herrando, M., Navarro, X. and Udina, E. (2015). Toxic effects of bortezomib on primary sensory neurons and Schwann cells of adult mice. Neurotox Res, 27, 430–440.

Alvarez-Castelao, B., Tom Dieck, S., Fusco, C. M., Donlin-Asp, P., Perez, J. D. and Schuman, E. M. (2020). The switch-like expression of heme-regulated kinase 1 mediates neuronal proteostasis following proteasome inhibition. Elife, 9.

Asghari Adib, E., Smithson, L. J. and Collins, C. A. (2018). An axonal stress response pathway: degenerative and regenerative signaling by DLK. Curr Opin Neurobiol, 53, 110–119.

Barlan, K. and Gelfand, V. I. (2017). Microtubule-Based Transport and the Distribution, Tethering, and Organization of Organelles. Cold Spring Harb Perspect Biol, 9.

Bauer, B., Martens, S. and Ferrari, L. (2023). Aggrephagy at a glance. J Cell Sci, 136.

Blasius, T. L., Reed, N., Slepchenko, B. M. and Verhey, K. J. (2013). Recycling of kinesin-1 motors by diffusion after transport. PLoS One, 8, e76081.

Blumenstock, S., Schulz-Trieglaff, E. K., Voelkl, K., Bolender, A. L., Lapios, P., Lindner, J., Hipp, M. S., Hartl, F. U., Klein, R. and Dudanova, I. (2021). Fluc-EGFP reporter mice reveal differential alterations of neuronal proteostasis in aging and disease. EMBO J, 40, e107260.

Cai, Q. and Ganesan, D. (2022). Regulation of neuronal autophagy and the implications in neurodegenerative diseases. Neurobiol Dis, 162, 105582.

Chiba, K., Kita, T., Anazawa, Y. and Niwa, S. (2023). Insight into the regulation of axonal transport from the study of KIF1A-associated neurological disorder. J Cell Sci, 136.

Chiba, K., Takahashi, H., Chen, M., Obinata, H., Arai, S., Hashimoto, K., Oda, T., McKenney, R. J. and Niwa, S. (2019). Disease-associated mutations hyperactivate KIF1A motility and anterograde axonal transport of synaptic vesicle precursors. Proc Natl Acad Sci U S A, 116, 18429–18434.

Chien, A., Shih, S. M., Bower, R., Tritschler, D., Porter, M. E. and Yildiz, A. (2017). Dynamics of the IFT machinery at the ciliary tip. Elife, 6.

Choi, W. H., Yun, Y., Park, S., Jeon, J. H., Lee, J., Lee, J. H., Yang, S. A., Kim, N. K., Jung, C. H., Kwon, Y. T., Han, D., Lim, S. M. and Lee, M. J. (2020). Aggresomal sequestration and STUB1-mediated ubiquitylation during mammalian proteaphagy of inhibited proteasomes. Proc Natl Acad Sci U S A, 117, 19190–19200.

Cohen, L. D., Zuchman, R., Sorokina, O., Muller, A., Dieterich, D. C., Armstrong, J. D., Ziv, T. and Ziv, N. E. (2013). Metabolic turnover of synaptic proteins: kinetics, interdependencies and implications for synaptic maintenance. PLoS One, 8, e63191.

Cong, D., Ren, J., Zhou, Y., Wang, S., Liang, J., Ding, M. and Feng, W. (2021). Motor domain-mediated autoinhibition dictates axonal transport by the kinesin UNC-104/KIF1A. PLoS Genet, 17, e1009940.

Conway, L., Wood, D., Tuzel, E. and Ross, J. L. (2012). Motor transport of self-assembled cargos in crowded environments. Proc Natl Acad Sci U S A, 109, 20814–20819.

Cross, J. A. and Dodding, M. P. (2019). Motor-cargo adaptors at the organelle-cytoskeleton interface. Curr Opin Cell Biol, 59, 16–23.

Di Domenico, F. and Lanzillotta, C. (2022). The disturbance of protein synthesis/degradation homeostasis is a common trait of age-related neurodegenerative disorders. Adv Protein Chem Struct Biol, 132, 49–87.

Ding, Q., Dimayuga, E., Markesbery, W. R. and Keller, J. N. (2006). Proteasome inhibition induces reversible impairments in protein synthesis. FASEB J, 20, 1055–1063.

Dorrbaum, A. R., Kochen, L., Langer, J. D. and Schuman, E. M. (2018). Local and global influences on protein turnover in neurons and glia. Elife, 7.

Engel, B. D., Ludington, W. B. and Marshall, W. F. (2009). Intraflagellar transport particle size scales inversely with flagellar length: revisiting the balance-point length control model. J Cell Biol, 187, 81–89.

Finkbeiner, S. (2020). The Autophagy Lysosomal Pathway and Neurodegeneration. Cold Spring Harb Perspect Biol, 12.

Frank, M., Citarella, C. G., Quinones, G. B. and Bentley, M. (2020). A novel labeling strategy reveals that myosin Va and myosin Vb bind the same dendritically polarized vesicle population. Traffic, 21, 689–701.

Gan, K. J., Akram, A., Blasius, T. L., Ramser, E. M., Budaitis, B. G., Gabrych, D. R., Verhey, K. J. and Silverman, M. A. (2020). GSK3beta Impairs KIF1A Transport in a Cellular Model of Alzheimer’s Disease but Does Not Regulate Motor Motility at S402. eNeuro, 7.

Geronimo-Olvera, C. and Massieu, L. (2019). Autophagy as a Homeostatic Mechanism in Response to Stress Conditions in the Central Nervous System. Mol Neurobiol, 56, 6594–6608.

Gupta, S.. T, M. R., Meng, F., Tidball, A., Akil, H., Watson, S., Parent, J. M. and Uhler, M. (2018). Fibroblast growth factor 2 regulates activity and gene expression of human post-mitotic excitatory neurons. J Neurochem, 145, 188–203.

Hakim, V., Cohen, L. D., Zuchman, R., Ziv, T. and Ziv, N. E. (2016). The effects of proteasomal inhibition on synaptic proteostasis. EMBO J, 35, 2238–2262.

Hammond, J. W., Cai, D., Blasius, T. L., Li, Z., Jiang, Y., Jih, G. T., Meyhofer, E. and Verhey, K. J. (2009). Mammalian Kinesin-3 motors are dimeric in vivo and move by processive motility upon release of autoinhibition. PLoS Biol, 7, e72.

Hendel, N. L., Thomson, M. and Marshall, W. F. (2018). Diffusion as a Ruler: Modeling Kinesin Diffusion as a Length Sensor for Intraflagellar Transport. Biophys J, 114, 663–674.

Hirokawa, N., Sato-Yoshitake, R., Kobayashi, N., Pfister, K. K., Bloom, G. S. and Brady, S. T. (1991). Kinesin associates with anterogradely transported membranous organelles in vivo. J Cell Biol, 114, 295–302.

Huang, H., Koyuncu, O. O. and Enquist, L. W. (2020). Pseudorabies Virus Infection Accelerates Degradation of the Kinesin-3 Motor KIF1A. J Virol, 94.

Hung, C. O. and Coleman, M. P. (2016). KIF1A mediates axonal transport of BACE1 and identification of independently moving cargoes in living SCG neurons. Traffic, 17, 1155–1167.

Hyer, M. L., Milhollen, M. A., Ciavarri, J., Fleming, P., Traore, T., Sappal, D., Huck, J., Shi, J., Gavin, J., Brownell, J., Yang, Y., Stringer, B., Griffin, R., Bruzzese, F., Soucy, T., Duffy, J., Rabino, C., Riceberg, J., Hoar, K., Lublinsky, A., Menon, S., Sintchak, M., Bump, N., Pulukuri, S. M., Langston, S., Tirrell, S., Kuranda, M., Veiby, P., Newcomb, J., Li, P., Wu, J. T., Powe, J., Dick, L. R., Greenspan, P., Galvin, K., Manfredi, M., Claiborne, C., Amidon, B. S. and Bence, N. F. (2018). A small-molecule inhibitor of the ubiquitin activating enzyme for cancer treatment. Nat Med, 24, 186–193.

Jacobson, C., Schnapp, B. and Banker, G. A. (2006). A change in the selective translocation of the Kinesin-1 motor domain marks the initial specification of the axon. Neuron, 49, 797–804.

Jin, Y. and Zheng, B. (2019). Multitasking: Dual Leucine Zipper-Bearing Kinases in Neuronal Development and Stress Management. Annu Rev Cell Dev Biol, 35, 501–521.

Kageyama, S., Sou, Y. S., Uemura, T., Kametaka, S., Saito, T., Ishimura, R., Kouno, T., Bedford, L., Mayer, R. J., Lee, M. S., Yamamoto, M., Waguri, S., Tanaka, K. and Komatsu, M. (2014). Proteasome dysfunction activates autophagy and the Keap1-Nrf2 pathway. J Biol Chem, 289, 24944–24955.

Kargbo-Hill, S. E. and Colon-Ramos, D. A. (2020). The Journey of the Synaptic Autophagosome: A Cell Biological Perspective. Neuron, 105, 961–973.

Kinger, S., Jagtap, Y. A., Kumar, P., Choudhary, A., Prasad, A., Prajapati, V. K., Kumar, A., Mehta, G. and Mishra, A. (2024). Proteostasis in neurodegenerative diseases. Adv Clin Chem, 121, 270–333.

Kumar, J., Choudhary, B. C., Metpally, R., Zheng, Q., Nonet, M. L., Ramanathan, S., Klopfenstein, D. R. and Koushika, S. P. (2010). The Caenorhabditis elegans Kinesin-3 motor UNC-104/KIF1A is degraded upon loss of specific binding to cargo. PLoS Genet, 6, e1001200.

Laser, H., Mack, T. G., Wagner, D. and Coleman, M. P. (2003). Proteasome inhibition arrests neurite outgrowth and causes “dying-back” degeneration in primary culture. J Neurosci Res, 74, 906–916.

Lazarevic, V., Schone, C., Heine, M., Gundelfinger, E. D. and Fejtova, A. (2011). Extensive remodeling of the presynaptic cytomatrix upon homeostatic adaptation to network activity silencing. J Neurosci, 31, 10189–10200.

Lee, J. R., Shin, H., Ko, J., Choi, J., Lee, H. and Kim, E. (2003). Characterization of the movement of the kinesin motor KIF1A in living cultured neurons. J Biol Chem, 278, 2624–2629.

Lee, K. D. and Hollenbeck, P. J. (1995). Phosphorylation of kinesin in vivo correlates with organelle association and neurite outgrowth. J Biol Chem, 270, 5600–5605.

Li, J., Cai, Z., Vaites, L. P., Shen, N., Mitchell, D. C., Huttlin, E. L., Paulo, J. A., Harry, B. L. and Gygi, S. P. (2021). Proteome-wide mapping of short-lived proteins in human cells. Mol Cell, 81, 4722–4735 e4725.

Li, J., Zhang, Y. V., Asghari Adib, E., Stanchev, D. T., Xiong, X., Klinedinst, S., Soppina, P., Jahn, T. R., Hume, R. I., Rasse, T. M. and Collins, C. A. (2017). Restraint of presynaptic protein levels by Wnd/DLK signaling mediates synaptic defects associated with the kinesin-3 motor Unc-104. Elife, 6.

Li, S., Wan, K. Y., Chen, W., Tao, H., Liang, X. and Pan, J. (2020). Functional exploration of heterotrimeric kinesin-II in IFT and ciliary length control in Chlamydomonas. Elife, 9.

Lienard, C., Pintart, A. and Bomont, P. (2024). Neuronal Autophagy: Regulations and Implications in Health and Disease. Cells, 13.

Ludington, W. B., Wemmer, K. A., Lechtreck, K. F., Witman, G. B. and Marshall, W. F. (2013). Avalanche-like behavior in ciliary import. Proc Natl Acad Sci U S A, 110, 3925–3930.

Ma, R., Hendel, N. L., Marshall, W. F. and Qin, H. (2020). Speed and Diffusion of Kinesin-2 Are Competing Limiting Factors in Flagellar Length-Control Model. Biophys J, 118, 2790–2800.

MacGillavry, H. D. and Hoogenraad, C. C. (2018). Membrane trafficking and cytoskeletal dynamics in neuronal function. Mol Cell Neurosci, 91, 1–2.

Mukherjee, T., Ramaglia, V., Abdel-Nour, M., Bianchi, A. A., Tsalikis, J., Chau, H. N., Kalia, S. K., Kalia, L. V., Chen, J. J., Arnoult, D., Gommerman, J. L., Philpott, D. J. and Girardin, S. E. (2021). The eIF2alpha kinase HRI triggers the autophagic clearance of cytosolic protein aggregates. J Biol Chem, 296, 100050.

Nam, T., Han, J. H., Devkota, S. and Lee, H. W. (2017). Emerging Paradigm of Crosstalk between Autophagy and the Ubiquitin-Proteasome System. Mol Cells, 40, 897–905.

Niwa, S., Lipton, D. M., Morikawa, M., Zhao, C., Hirokawa, N., Lu, H. and Shen, K. (2016). Autoinhibition of a Neuronal Kinesin UNC-104/KIF1A Regulates the Size and Density of Synapses. Cell Rep, 16, 2129–2141.

Oettinger, D. and Yamamoto, A. (2025). Autophagy Dysfunction and Neurodegeneration: Where Does It Go Wrong? J Mol Biol, 437, 169219.

Okada, Y., Yamazaki, H., Sekine-Aizawa, Y. and Hirokawa, N. (1995). The neuron-specific kinesin superfamily protein KIF1A is a unique monomeric motor for anterograde axonal transport of synaptic vesicle precursors. Cell, 81, 769–780.

Okita, K., Matsumura, Y., Sato, Y., Okada, A., Morizane, A., Okamoto, S., Hong, H., Nakagawa, M., Tanabe, K., Tezuka, K., Shibata, T., Kunisada, T., Takahashi, M., Takahashi, J., Saji, H. and Yamanaka, S. (2011). A more efficient method to generate integration-free human iPS cells. Nat Methods, 8, 409–412.

Ross, A. B., Langer, J. D. and Jovanovic, M. (2021). Proteome Turnover in the Spotlight: Approaches, Applications, and Perspectives. Mol Cell Proteomics, 20, 100016.

Sabharwal, V. and Koushika, S. P. (2019). Crowd Control: Effects of Physical Crowding on Cargo Movement in Healthy and Diseased Neurons. Front Cell Neurosci, 13, 470.

Selvaraj, C., Vijayalakshmi, P., Desai, D. and Manoharan, J. (2025). Proteostasis imbalance: Unraveling protein aggregation in neurodegenerative diseases and emerging therapeutic strategies. Adv Protein Chem Struct Biol, 146, 1–34.

Shen, D., Coleman, J., Chan, E., Nicholson, T. P., Dai, L., Sheppard, P. W. and Patton, W. F. (2011). Novel cell- and tissue-based assays for detecting misfolded and aggregated protein accumulation within aggresomes and inclusion bodies. Cell Biochem Biophys, 60, 173–185.

Shen, Y. F., Tang, Y., Zhang, X. J., Huang, K. X. and Le, W. D. (2013). Adaptive changes in autophagy after UPS impairment in Parkinson’s disease. Acta Pharmacol Sin, 34, 667–673.

Shin, J. E., Miller, B. R., Babetto, E., Cho, Y., Sasaki, Y., Qayum, S., Russler, E. V., Cavalli, V., Milbrandt, J. and DiAntonio, A. (2012). SCG10 is a JNK target in the axonal degeneration pathway. Proc Natl Acad Sci U S A, 109, E3696–3705.

Sidibe, D. K., Vogel, M. C. and Maday, S. (2022). Organization of the autophagy pathway in neurons. Curr Opin Neurobiol, 75, 102554.

Sleigh, J. N., Rossor, A. M., Fellows, A. D., Tosolini, A. P. and Schiavo, G. (2019). Axonal transport and neurological disease. Nat Rev Neurol, 15, 691–703.

Smith, G., Sweeney, S. T., O’Kane, C. J. and Prokop, A. (2023). How neurons maintain their axons long-term: an integrated view of axon biology and pathology. Front Neurosci, 17, 1236815.

Smith, T. P., Sahoo, P. K., Kar, A. N. and Twiss, J. L. (2020). Intra-axonal mechanisms driving axon regeneration. Brain Res, 1740, 146864.

Staff, N. P., Podratz, J. L., Grassner, L., Bader, M., Paz, J., Knight, A. M., Loprinzi, C. L., Trushina, E. and Windebank, A. J. (2013). Bortezomib alters microtubule polymerization and axonal transport in rat dorsal root ganglion neurons. Neurotoxicology, 39, 124–131.

Stucchi, R., Plucinska, G., Hummel, J. J. A., Zahavi, E. E., Guerra San Juan, I., Klykov, O., Scheltema, R. A., Altelaar, A. F. M. and Hoogenraad, C. C. (2018). Regulation of KIF1A-Driven Dense Core Vesicle Transport: Ca(2+)/CaM Controls DCV Binding and Liprin-alpha/TANC2 Recruits DCVs to Postsynaptic Sites. Cell Rep, 24, 685–700.

Xiong, G. J. and Sheng, Z. H. (2024). Presynaptic perspective: Axonal transport defects in neurodevelopmental disorders. J Cell Biol, 223.

Xiong, X., Wang, X., Ewanek, R., Bhat, P., Diantonio, A. and Collins, C. A. (2010). Protein turnover of the Wallenda/DLK kinase regulates a retrograde response to axonal injury. J Cell Biol, 191, 211–223.

Yildiz, A. (2025). Mechanism and regulation of kinesin motors. Nat Rev Mol Cell Biol, 26, 86–103.

Zhang, L., Ebenezer, P. J., Dasuri, K., Bruce-Keller, A. J., Liu, Y. and Keller, J. N. (2009). Proteasome inhibition modulates kinase activation in neural cells: relevance to ubiquitination, ribosomes, and survival. J Neurosci Res, 87, 3231–3238.

